# Targeting high-density aromatic peptides to cardiolipin optimizes the mitochondrial membrane potential

**DOI:** 10.1101/2023.09.28.560027

**Authors:** Alexander Birk, Shaoyi Liu, Virginia Garcia-Marin, Margaret A. MacNeil

## Abstract

The mitochondrial membrane potential (ΔΨm) is created by the accumulation of protons on an outer leaflet of the inner mitochondrial membrane and drives the synthesis of most cellular ATP, which is essential for cellular bioenergetics and survival. The ΔΨm also facilitates the electrogenic transport of cations, such as Ca^2+^, and regulates generation of reactive oxygen species, which serves as a powerful bioenergetic and stress-signaling regulator. Proton trapping on the outer leaflet of the inner mitochondrial membrane of mitochondrial cristae could be controlled by cardiolipin when the local pH is above 8. However, there is presently no technology that effectively targets strong bases to cardiolipin.

We have developed a novel, high-density aromatic peptide (HDAP2) to preserve a proton gradient-driven potential in mitochondria by increasing proton trapping on cardiolipin (CL). HDAP2-induced formation of cardiolipin-HDAP2 complexes accumulated positive charges at the head of CL. The HDAP2-CL vesicles could accumulate the mitochondrial transmembrane potential probe, Tetramethylrhodamine (TMRM). This potential could be uncoupled with Carbonyl cyanide m-chlorophenylhydrazone (CCCP) and Dinitrophenol (DNP), indicating that an interaction of HDAP2 with CL could support a proton gradient-driven transmembrane potential.

We demonstrated that this novel, water-soluble peptide is cell-permeable, targets mitochondria without causing cell toxicity, and promotes cell survival during serum starvation. Importantly, the HDAP2-cardiolipin complex-mediated optimization of the proton gradient was supported by the ability of HDAP2 to prevent CCCP-mediated mitochondrial depolarization in ARPE-19 cells in a dose-dependent manner. Based on its mechanism of action, HDAP2 could promote cellular homeostasis, which would have broad clinical applicability for the prevention, recovery and reversal of many acute and chronic disease conditions, such as neurodegeneration, ischemia– reperfusion injury, and inflammation.

## INTRODUCTION

Mitochondria play a pivotal role in the physiology and pathology of many acute and chronic disease conditions, such as cancer (1, 2), neurodegeneration, ischemia– reperfusion injury, stroke, inflammation, and a variety of infections (3, 4). Stability and careful regulation of the mitochondrial membrane potential (ΔΨm) is required for supporting cellular bioenergetics, normal metabolism, and normal cell functioning, by promoting cellular ATP synthesis and maintaining Ca^2+^ and Na^+^ homeostasis (5, 6). Myocardial infarction (7) and stroke (8) can be initiated by a loss of ΔΨm and dissipation of the ΔΨm precedes mitochondria-mediated apoptosis (9-12).

Generation of the ΔΨm is accomplished by promoting coupling of the electron transport chain with proton transfer into the cristae-defined intermembrane space (6). Mitochondrial depolarization is induced by inhibition of the electron transport chain, promoting proton leakage from the mitochondrial inner membrane space into its matrix, stimulating mitochondrial uncoupling proteins, using uncouplers, and transporting positively charged molecules into the mitochondrial matrix (6).Trapping protons within the outer leaflet of the inner mitochondrial membrane, where both ATP synthase and cardiolipin are densely concentrated (13, 14), could be a way to preserve the ΔΨm. The process is thought to be controlled by cardiolipin when the local pH is above 8 (15, 16), suggesting that delivery of strong bases to the cardiolipin head could potentially promote proton trapping and support the ΔΨm. However, there is presently no technology that effectively targets strong bases to cardiolipin.

Introducing electroconductive materials into the mitochondrial membrane could have additional benefits of optimizing the electron transport chain and the ΔΨm. Past attempts at using Phe-Phe for this purpose, a dipeptide with known conductive characteristics, was toxic to cells (17-21). In this communication, we describe a novel peptide that localizes aromatic Phe next to cationic Arginine, which effectively brackets the high-density aromatic Phe-Phe dipeptide rendering it nontoxic to cells. Additionally, we will demonstrate that this high-density aromatic peptide (HDAP2) interacts selectively with cardiolipin, promotes a proton-driven transmembrane potential in HDAP2-cardiolipin vesicles, targets mitochondria in live cells, protects against cell toxicity during serum-starvation in MDBK cells, and prevents CCCP-mediated loss of ΔΨm in ARPE-19 cells.

## METHODOLOGY

### 1) Chemicals

HDAP2 (>96.5 % pure) was synthesized by GeneScript (Piscataway, NJ). Other chemicals, including 1-palmitoyl-2-oleoyl-sn-glycero-3-phosphocholine (POPC), cardiolipin from bovine heart (CL, consisting primarily of tetralinoleoyl CL), reagents and assay kits were purchased from Sigma Aldrich (St. Louis, MO).

### 2) Preparation of Liposomes

Lipids in chloroform were combined in 12×75 mm glass tubes with either 150 μM CL: 150 μM POPC or 300 μM POPC. The solvent was allowed to slowly evaporate, which resulted in a lipid film that could be rehydrated in an aqueous solution of 10 mM HEPES, pH 7.4. The resulting multilamellar vesicles were vortexed lightly and sized into small unilamellar vesicles by heated bath sonication for 25 min. All liposomes were cooled to ambient temperature before use.

### 3) Interaction of HDAP2 Peptide with Phospholipids

We determined the ability of HDAP2 to displace nonyl acridine orange (NAO), a fluoroprobe known to selectively bind CL (22, 23). Interaction of HDAP2 with phospholipids was examined by measuring changes in the fluorescence spectra of 3 μM NAO (Molecular Devices, Sunnyvale, CA)) with excitation at 480 nm. CL and POPC in liposomes were use at 30 μM. Dose response curves were done with 30 μM CL, HDAP2 (0, 2.5, 5, 10, 20, and 30 μM), and 3 μM of NAO (ex/em at 480 nm/520 nm). All experiments were done in 10 mM Hepes (pH 7.4) to optimize electrostatic interactions of peptides with phospholipids. The different solvents used to dissolve the various phospholipids (chloroform, methanol, and ethanol) had a negligible effect on the fluorescence spectra.

### 4) Interactions between HDAP2 and Phospholipid liposomes measured by Nile Red fluorescence

The hydrophobicity of HDAP2 and phospholipids was examined by changes in the spectral fluorescence shift and intensity of Nile Red (NR, 10 μM). NR increases its intensity of fluorescence when combined with the hydrophobic environments of acyl chains of phospholipids, and produces a blue shift when hydrophobicity is increased (24). All experiments were done with 30 μM of either CL or POPC and 30 μM of HDAP2 in 10 mM Hepes (pH 7.4) to optimize the electrostatic interactions of peptides with phospholipids. The solvents used to dissolve the various phospholipids (chloroform, methanol, and ethanol) had negligible effects on the fluorescence spectra.

### 4) Preparation and Visualizing HDAP2-CL vesicles with Nile Red

To visualize HDAP2-CL vesicles with NR, 30 μM of CL or POPC were mixed with 100 μM of HDAP2 in 10 mM Hepes pH 7.4. The mixture was sonicated for 30 min, transferred to 35 mm cell culture wells, and allowed to settle for 24 hours. The next day, 10 μM of NR was added for 15 min at room temperature and then gently washed three times to remove excess NR. Samples with no peptide or phospholipids were used as controls. Fluorescence staining was observed by using a Nikon Eclipse 50i. The intensity of NR fluorescence was measured in each field, using the “Analyze” feature in ImageJ to compute the mean pixel intensity (in arbitrary fluorescence units) from the images. Differences among groups were compared by one-way ANOVA. Post hoc analyses were carried out using Tukey’s multiple comparisons test.

### 5) Labeling HDAP2-CL vesicles with TMRM

After forming HDAP2-phospholipids vesicles, 100 nM of TMRM was added for 15 min at room temperature and then gently washed three times to remove excess TMRM. The sample with no peptide or CL was used as control. For testing proton-gradient uncouplers, 100 μM CCCP or 1 mM DNP was added together with TMRM and imaged and analyzed as described for NR.

### 6) Intracellular Localization of HDAP2

Intracellular targeting of HDAP2 was determined by incubating MDBK cells (ATCC-CCL22) that were grown in biotin-free media for 72 hours, with 10 μM of N-biotinylated HDAP2 for 1 hour. Cells were then fixed with 4% paraformaldehyde, permeabilized with 0.1% Triton X-100, and treated with streptavidin conjugated with Alexa Fluor 488 (SA) for 30 minutes at room temperature. The Hoechst 33342 probe was used to label the nucleus to determine cellular localization and relative distributions of HDAP2 and other cellular probes. Fluorescence staining was observed with a Nikon Eclipse 50i microscope (20× objective). To determine mitochondrial localization of HDAP2, the same cells were also stained with 10 nM of MitoTracker red CMXRos, before fixation. SA fluorescence was normalized to Hoechst fluorescence to account for differences in cell number in each field. The intensity of SA fluorescence was measured in each field, using the “Analyze” feature in ImageJ to compute the mean pixel intensity (in arbitrary fluorescence units) from the images. Differences among groups were compared by one-way ANOVA. Post hoc analyses were carried out using Tukey’s multiple comparisons test.

### 7) Effects of peptides on cell survival in serum-free media

Cell survival in serum starvation conditions was studied in the presence and absence of HDAP2 using MDBK cells. Cells were plated at a density of 1–2 × 10^3^ cells/well in 96 well plates in DMEM/10% FBS media. The following day, cells were washed and the medium replaced with DMEM serum-free media in the presence or absence of different concentrations of HDAP2 and incubated for 5 days. Cell viability was assessed using the Resaruzin (Alamar blue) indicator dye (25). Quantitative analysis of dye conversion was measured using a fluorescent plate reader (ex/em = 550/580) and viability was expressed as fold increase over of untreated cells.

### 8) Effects of peptides on mitochondrial potential in the presence of Carbonyl cyanide m-chlorophenylhydrazone (CCCP)

ARPE-19 cells (ATTC-CRL-2302) (5 × 10^4^ cells) were seeded in 35 mm glass dishes in DMEM/F12 medium with 10% FBS for 24 hr. On the day of the experiment, cells were cultured in DMEM/F12 in the presence of different concentrations of HDAP2 for 60 min (Fig 4B). Then, cells were incubated with 10 μM CCCP in the presence of 5 nM TMRM, 100 nM MitoView Green, and 10 μg/ml Hoechst 33342 for 15 min at 37°C. Fluorescent images were immediately obtained with a Nikon Eclipse 50i fluorescence microscope (60× objective) using FITC (MitoView Green), Texas Red (TMRM), and DAPI (Hoechst) filters. Images were collected from three independent experiments, using 4–6 random fields for each treatment group, and imaged and analyzed as described for NR.

## RESULTS

Inserting electroconductive materials, such as a Phe-Phe dipeptide motif, into the mitochondrial membrane could potentially optimize the electron transport chain and the ΔΨm. However, past attempts at using Phe-Phe for this purpose were toxic to cells (17-21). Our novel peptide HDAP2 is organized with aromatic amino acids (Phe) next to positively charged amino acids (Arg), which create a structural motif that promotes interactions with cardiolipin (26). We wanted to test whether peptides with this general motif would interact with cardiolipin and promote the ΔΨm.

We tested the selectivity of HDAP2 for CL by demonstrating that HDAP2 displaced binding of a known CL-selective fluoroprobe, nonyl acridine orange (NAO) (22, 23), from POPC-CL liposomes (Fig 1 A-C). When NAO was mixed with CL, the intensity of its fluorescence increased by about 3-fold, compared to the intensity of fluorescence when NAO was mixed with POPC or HDAP2 alone (Fig 1 A and B). When also combined with CL, HDAP2 decreased the intensity of NAO fluorescence to background levels, and did not interfere with the interaction between NAO and the POPC liposomes (Fig 1 A and B), suggesting the selectivity of HDAP2 to CL. In addition, we determined that HDAP2 outcompeted NAO for binding with CL, with EC50 at a ratio of about one peptide to two CL molecules, suggesting that HDAP2 can bind two CL simultaneously. We also confirmed the selective interaction of HDAP2 with CL by using the dye Nile Red (NR), which selectively binds to acyl chains of phospholipids by producing a blue shift in its emission (24). In POPC-CL liposomes, HDAP2 promoted a blue shift of NR from 638 nm to 618 nm (Fig 1D middle panel), but no shift could be detected when HDAP2 was added to POPC liposomes (Fig 1D, bottom panel), confirming the selectivity of HDAP2 for CL.

**Figure 1.**
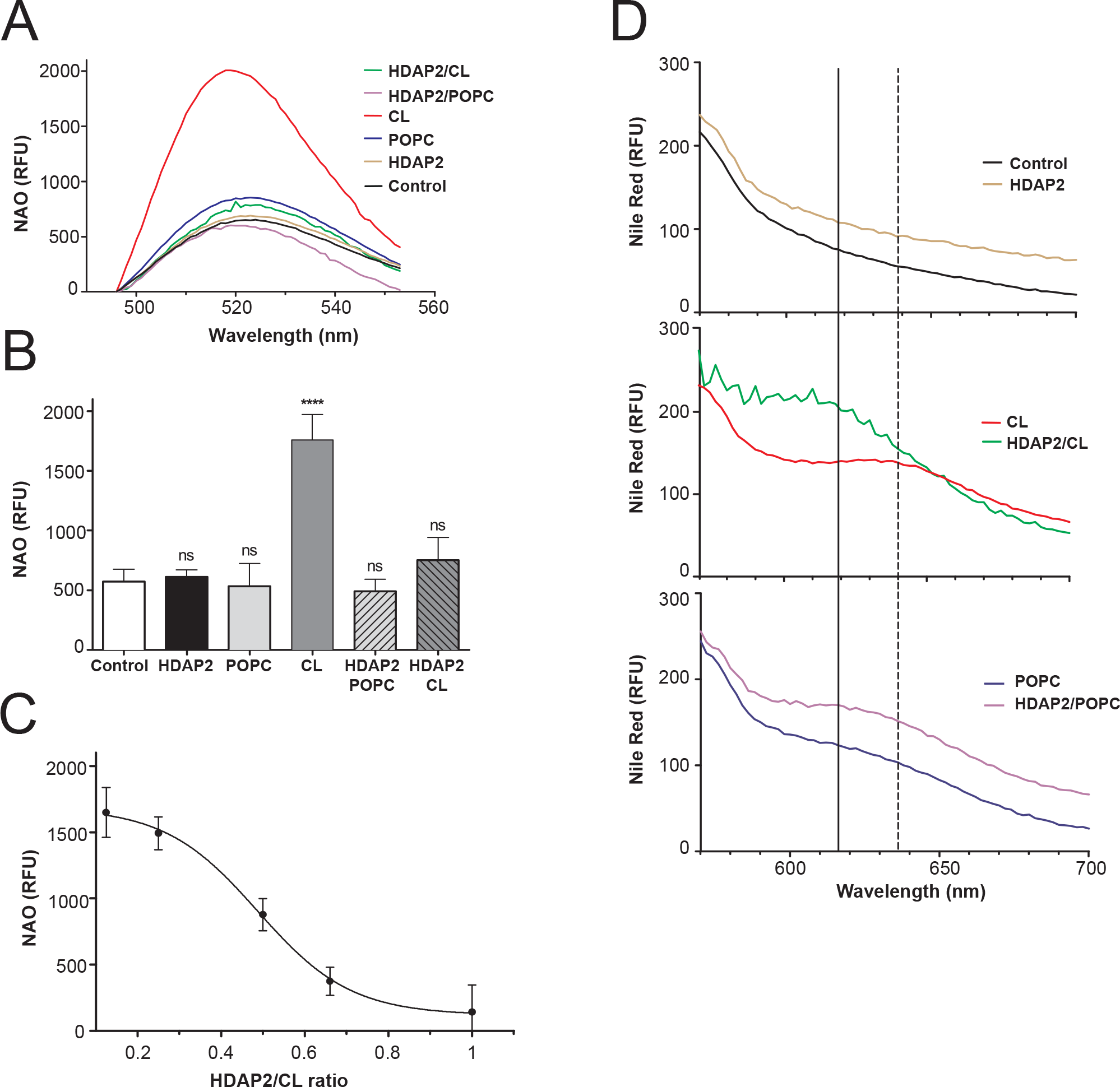
(A) Representative fluorescence emission spectra of NAO (3 μM, λex=480 nm) in the presence of different POPC (60 μM) and POPC-CL liposomes (POPC 30 μM and CL 30 μM) and 30 μM HDAP2. (B) Quantitative analysis of the effect of different phospholipids and HDAP2 on NAO fluorescence. (C) HDAP2 decreases the intensity of NAO fluorescence with POPC-CL liposomes in a dose-dependent manner. (D) Blue shifts in NR emission maximum (λmax) were observed in POPC-CL liposomes (dotted vertical line, 638 nm) in the presence of HDAP2 (solid vertical line, 618 nm), but not in POPC liposomes or HDAP2 alone. Error bars represent SEM (n=6) ***P < 0.001.

The blue shift observed from the NR experiments suggests that HDAP2 increases the hydrophobicity of the acyl chain in CL and would also indicate that d-Arg is likely to be at the CL head in the lipid-water interphase. Arg would provide positive charge to an HDAP2-CL complex, which can be bound to negatively charged surfaces. To test this hypothesis, we combined CL and HDAP2, without additional POPC. Surprisingly, we observed that HDAP2-CL vesicular structures adhered to the negative surface of the cell culture wells and were stained with NR, unlike blank wells that were unstained (Fig 2 A right panel and B). Vesicular structures or NR staining were not observed when HDAP2 was mixed with POPC in the blank wells (Fig 2 A and B) or when added to untreated wells (data not shown), further demonstrating selectivity of HDAP2 for CL.

**Figure 2.**
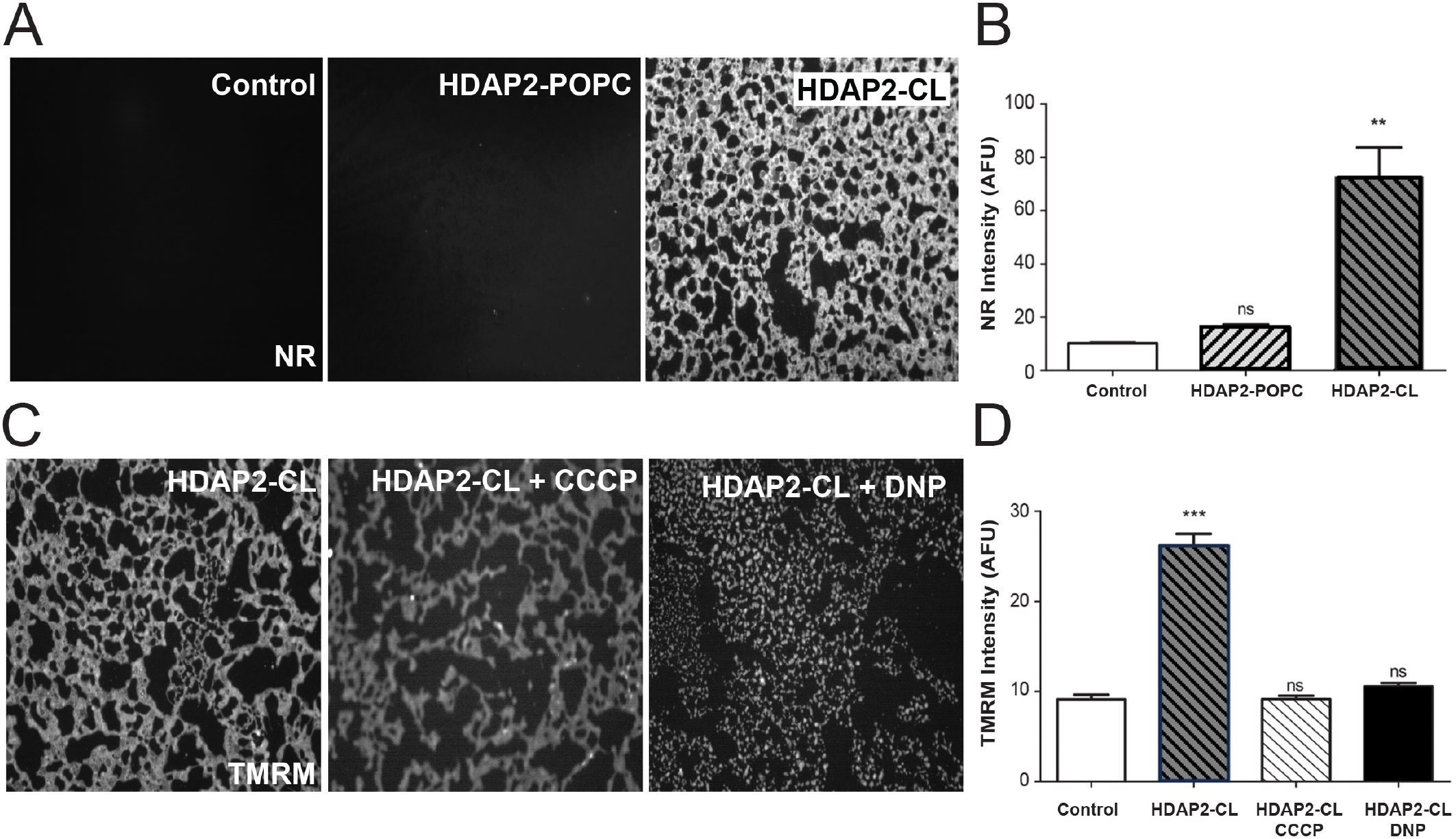
(A) Detection of POPC-HDAP2 (middle panel) and CL-HDAP2 vesicles on the surface of cell culture plates-treated with 100nM NR. (B) Intensity of NR labeling of with HDAP2-POPC and HDAP2-CL vesicles. Error bars represent SEM (n=6) **P < 0.01. (C) Representative images of CL-HDAP2 vesicles labeled with 100nM of TMRM alone (left panel) and in the presence of proton gradient uncouplers 100 μM of CCCP (middle panel) and 1 mM DNP (right panel). (D) Quantitative analysis of TMRM-dependent labeling of CL-HDAP2 vesicles in the presence of CCCP and DNP. Error bars represent SEM (n=6) ***P < 0.001.

Concentration of alkaline Arg on the surface of HDAP2-CL vesicles could potentially trap protons on CL to create polarization of a HDAP2-CL membrane. We used TMRM, a proton-gradient transmembrane potential probe, commonly used in mitochondria research, to show that HDAP2-CL vesicles were stained (Fig 2 C left panel and D), indicating a transmembrane proton potential across the membrane of the vesicles. To confirm that the potential across the membrane was due to proton trapping of the CL-HDAP2 surface, we measured significant decreases in TMRM staining in the presence of proton ionophores, Carbonyl cyanide m-chlorophenylhydrazone (CCCP) and Dinitrophenol (DNP) (Fig 2 C (middle and right panels and D). Our data supports the idea that HDAP2 traps protons on HDAP2-CL complex and generates a transmembrane proton potential.

To determine the activity of HDAP2 on mitochondria *in vivo*, we first demonstrated that it could be taken up into MDBK cells. Cells dramatically deplete mitochondrial biotin after 3 days of incubation in serum-free DMEM (27), yet biotinylated HDAP2 was taken up into the serum-starved cells after just 1 hour of exposure with the peptide (Fig 3 A right panel and B). HDAP2 was distributed in the perinuclear area, as indicated by streptavidin staining around nuclei (stained by Hoechst 33342) and entered the cell about 6 times faster than observed for biocytin (Fig 3 A middle panel and Fig B), suggesting that mechanisms used to take up HDAP2 into cells are likely to be peptide structure dependent. To determine mitochondria-targeting of HDAP2, we demonstrated that HDAP2 is colocalized with the selective mitochondrial marker, MitoTracker red CMXRos (Fig 3C) and found that it promoted cell survival of MDBK cells in serum-free conditions with EC50 of about 100 nM (Fig 3 D).

**Figure 3.**
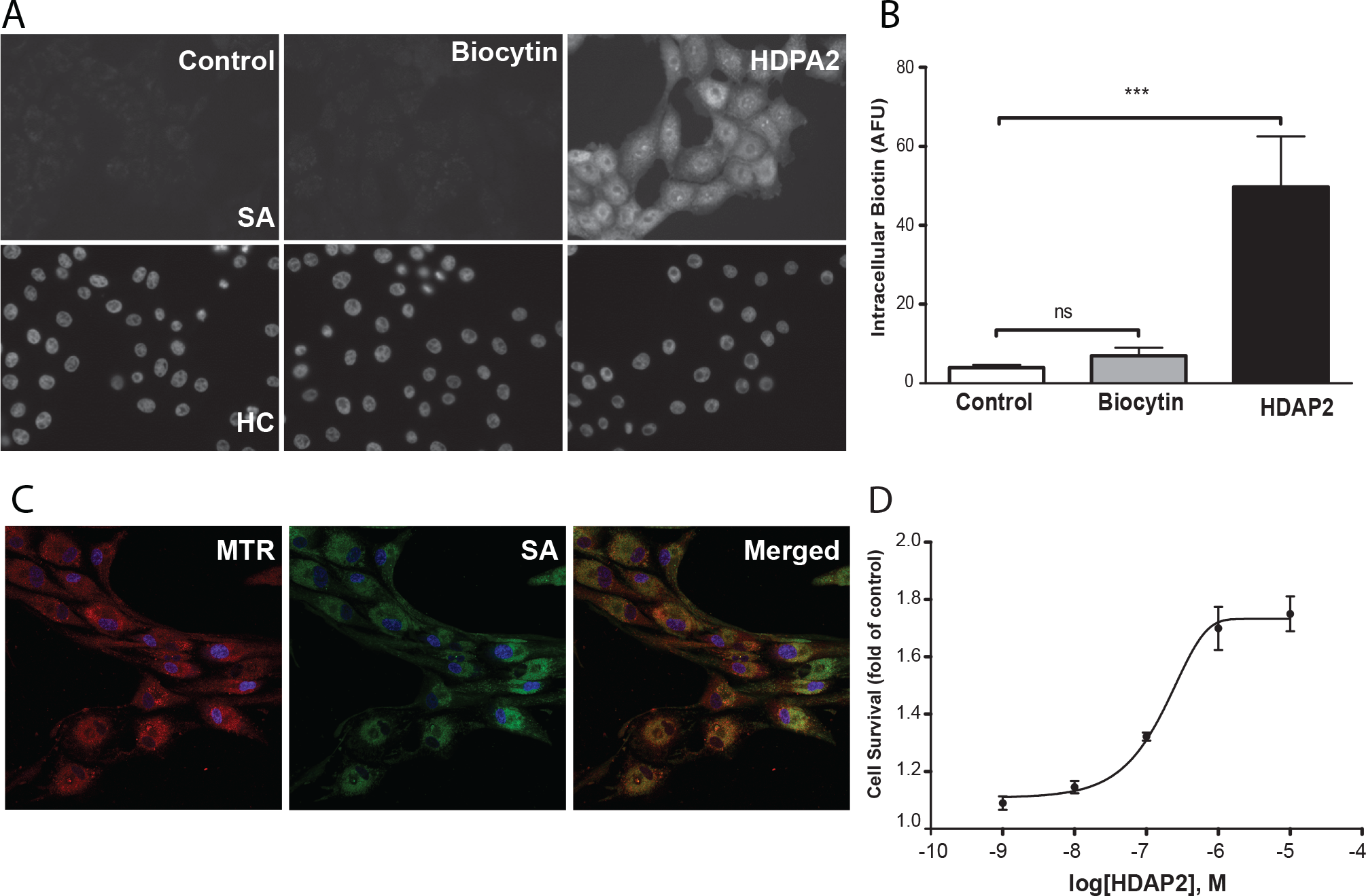
(A) Representative images of MDBK cells incubated with and streptavidin AlexaFluor 488 to demonstrate that 10 μM of N-biotinylated HDAP2 readily loads into cells (right panel), compared with 10 μM of biocytin (middle panel). (B) Quantitative analysis of HDAP2 and biocytin uptake in MDBK cells. Error bars represent SEM (n=5) ***P < 0.001. (C) Cultured MDBK cells labeled with streptavidin AF488 to identify HDAP2 (green) and Mitotracker Red CMXRos (red) to label mitochondria. Co-localization is indicated by the yellow labeling. (D) HDAP2 promotes survival of serum-starved MDBK cells in a dose-dependent manner, with EC50 of 100 nM.

**Figure 4.**
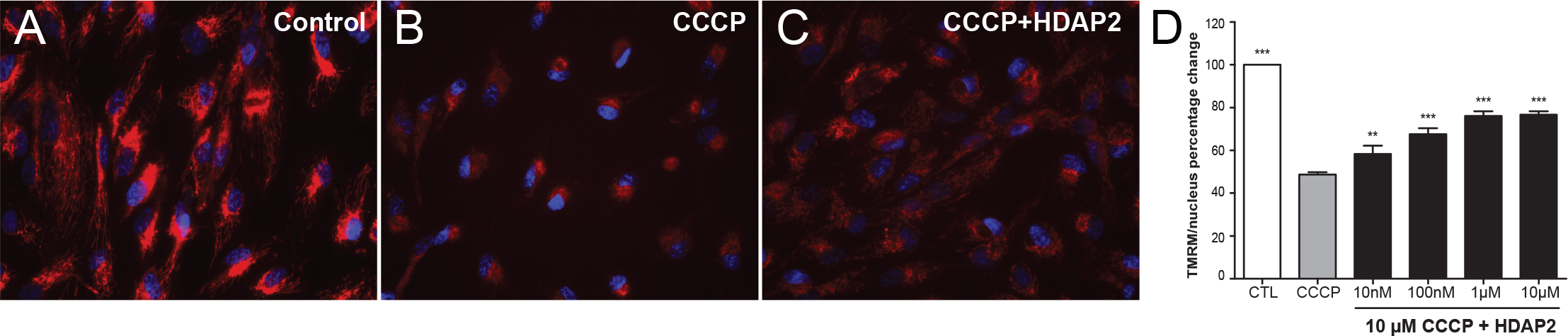
(A) ARPE-19 cells incubated with TMRM to demonstrate changes in the mitochondrial membrane potential (red). When cells were incubated with 10 μM CCCP (middle), TMRM staining was virtually extinguished), but showed increased fluorescence when 1 μM HDAP2 was added in addition to 10 μM CCCP. (B) HDAP2 prevented CCCP-mediated decreases in the mitochondrial membrane potential in a dose-dependent manner. Error bars represent SEM (n=5) ***P < 0.001.

Finally, we hypothesized that an optimization of proton trapping within the HDAP2-CL complex on the inner mitochondrial membrane (IMM) has a potential to provide some resistance to the mitochondrial uncouplers. To test this hypothesis, we characterized the effect of HDAP2 on the ΔΨm decrease in the presence of known mitochondrial uncoupler CCCP in ARPE-19 cells. These cells were specifically chosen for their robust mitochondria and their ability to survive even when exposed to mitochondrial stressors. This allowed us to study rapid changes in ΔΨm without inducing cell toxicity from overlapping events. We demonstrated that even 10 μM of CCCP dropped the ΔΨm to 20% of control cells (Fig 4 and B). Then, we determined that HDAP2 prevented mitochondrial depolarization in a dose-dependent manner when incubated together with 10 μM of CCCP (Fig 4 A and B), suggesting that HDAP2-mediated proton trapping might be a novel recoupling mechanism to preserve the ΔΨm.

## DISCUSSION

In this communication, we tested the idea of using the electroconductive Phe-Phe motif to bring positively charged Arg to cardiolipin and promote proton trapping on cardiolipin. However, Phe-Phe self-ensembles into fibrils, and when combined with positively charged triphenylphosphonium, it can accumulate in the mitochondrial matrix, causing mitochondrial depolarization and cellular toxicity. On the other hand, alternating aromatic-cationic motif binds to cardiolipin on the inner mitochondrial membrane, but it was never demonstrated to have direct effect on mitochondrial potential or proton trapping (26-29).

We have demonstrated that combining Phe-Arg and Phe-Phe motifs in HDAP2 promotes an interaction of HDAP2 with cardiolipin. As it was previously demonstrated for the cardiolipin-binding peptide SS-31(27), HDAP2 outcompeted an interaction of the selective cardiolipin probe, NAO, with CL in a dose-dependent manner. In addition, we demonstrated that HDAP2 binds to CL in a 1:2 ratio, which indicates stronger and more specific binding than found for SS peptides and CL, which have a 1:1 ratio (27). Phe is thought to penetrate deeply into the cardiolipin-containing phospholipid bilayer (28), which would increase hydrophobicity of acyl groups in the phospholipid bilayer. To support this hypothesis, we demonstrated that HDAP selectively promotes a NR blue shift in CL-containing POPC liposomes, without affecting POPC liposomes on their own.

This interaction of Phe with acyl chains of CL suggests that the two Arg within HDAP are likely to localize near the head group of CL on the phospholipid-soluble interphase and creates positive charge at the CL head. We wanted to test whether we could increase these positive charges by using a combination of HDAP2 with CL, without POPC. Surprisingly, HDAP2 formed vesicular structures with CL that were visualized with NR. These structures attached to the negative surface of cell culture-treated plates and could not be removed by washing with Hepes buffer. In contrast, POPC, which also forms liposomes, did not remain on the cell culture plates after washing, which supports our hypothesis. It is possible that HDAP2 interacts with acyl chains of CL in an inverted micele, but acyl chains of CL are similar to acyl chains of POPC, suggesting that the selective interaction of HDAP2 with CL is likely to be on the level of CL head group. This arrangement has also been described for other CL-targeted peptides, but work will be required to understand the detailed arrangement of HDAP2-CL vesicular structures.

Accumulation of Arg at the head group of CL is likely to increase the local pH around the hydroxyl group, which would promote proton-trapping conditions for CL. This explains why TMRM accumulated in HDAP2-CL vesicles. Accumulation of TMRM was inhibited by known uncouplers, CCCP and DNP, which supports the idea that HDAP2-CL vesicles create transmembrane proton gradients. Since 10 mM Hepes at pH 7.4 is likely to neutralize pH change, due to free Arg in the bulk solvent phase, HDAP2 possibly promotes accumulation of protons on the phospholipid-soluble interphase and preserves proton-mediated membrane potential, which further supports the idea of proton trapping in HDAP2-CL complexes.

To test whether HDAP2 can promote potential in mitochondria *in vivo*, we first demonstrated whether HDAP2 could be taken up into cells. After treating serum-starved cells with N-terminus biotinylated HDAP2 for one hour, we found a five-fold increase in streptavidin labeling compared to control cells or to cells incubated with an equimolar concentration of biocytin. The rapid loading of biotinylated HDAP2 compared to biocytin, suggests that HDAP2 cellular uptake was driven by the structure of the peptide, which is also the case for other CL-targeted peptides (26-30). In addition, biotinylated HDAP2 co-localized with the specific mitochondria markers, MitoTracker red CMXRos (Fig 3C) and Mitotracker green (data not shown), indicating that HDAP2 targets mitochondria.

Since previous studies have shown that targeting the Phe-Phe dipeptide to mitochondria is cytotoxic, it was important to test the toxicity of HDAP2 in cells. We demonstrated that incubation of 100 μM of HDAP2 with MDBK and ARPE-19 cells had no effect on cell growth (data not shown). HDAP2 also prevented serum starvation-mediated MDBK cell death in a dose-dependent manner, with EC50 of about 100 nM. It is important to point out that since HDAP2 has an overall positive charge at physiological pH, the delivery of these positive charges into the mitochondrial matrix is likely to lower the mitochondrial potential and induce cell death (31). The fact that HDAP2 is not cytotoxic suggests that HDAP2 does not enter the mitochondrial matrix.

CL is concentrated in the inner mitochondrial membrane where an interaction of HDAP2 and CL could produce proton trapping and protect the mitochondrial membrane potential against the uncoupling effects of CCCP. Since HDAP2 protects the membrane potential against CCCP activity, our results suggest that the peptide interacts with CL on the outer leaflet of the inner mitochondrial membrane where CCCP typically interacts with protons to transfer them into the matrix. Anti-CCCP activity of HDAP2 can also be explained by the trapping of protons on the outer leaflet of the inner mitochondrial membrane. The mechanism used by the HDAP2-CL complex to trap protons is novel and requires further investigation (32-34).

Our communication demonstrates that HDAP2, a novel CL-targeted peptide, promotes proton trapping within the HDAP2-CL complex and preserves the mitochondrial membrane potential in pathological conditions. Based on its mechanism of action, HDAP2 could promote mitochondrial ATP synthesis, support intracellular and mitochondrial Na+ and Ca2+ homeostasis, and prevent mitochondria-mediated apoptosis. These attributes give HDAP2 broad clinical applicability for the prevention, recovery and reversal of many acute and chronic disease conditions, such as neurodegeneration, ischemia–reperfusion injury, and inflammation.

